## Supplementary material for "DCs targeted therapy expands CD8 T cell responses to bona-fide neoantigens in lung tumors": Supp_file_Lopez_et_al_2024.pdf

**SUPPLEMENTARY DATA FILE**

**This file contains Supplementary Figures S1 to S6**

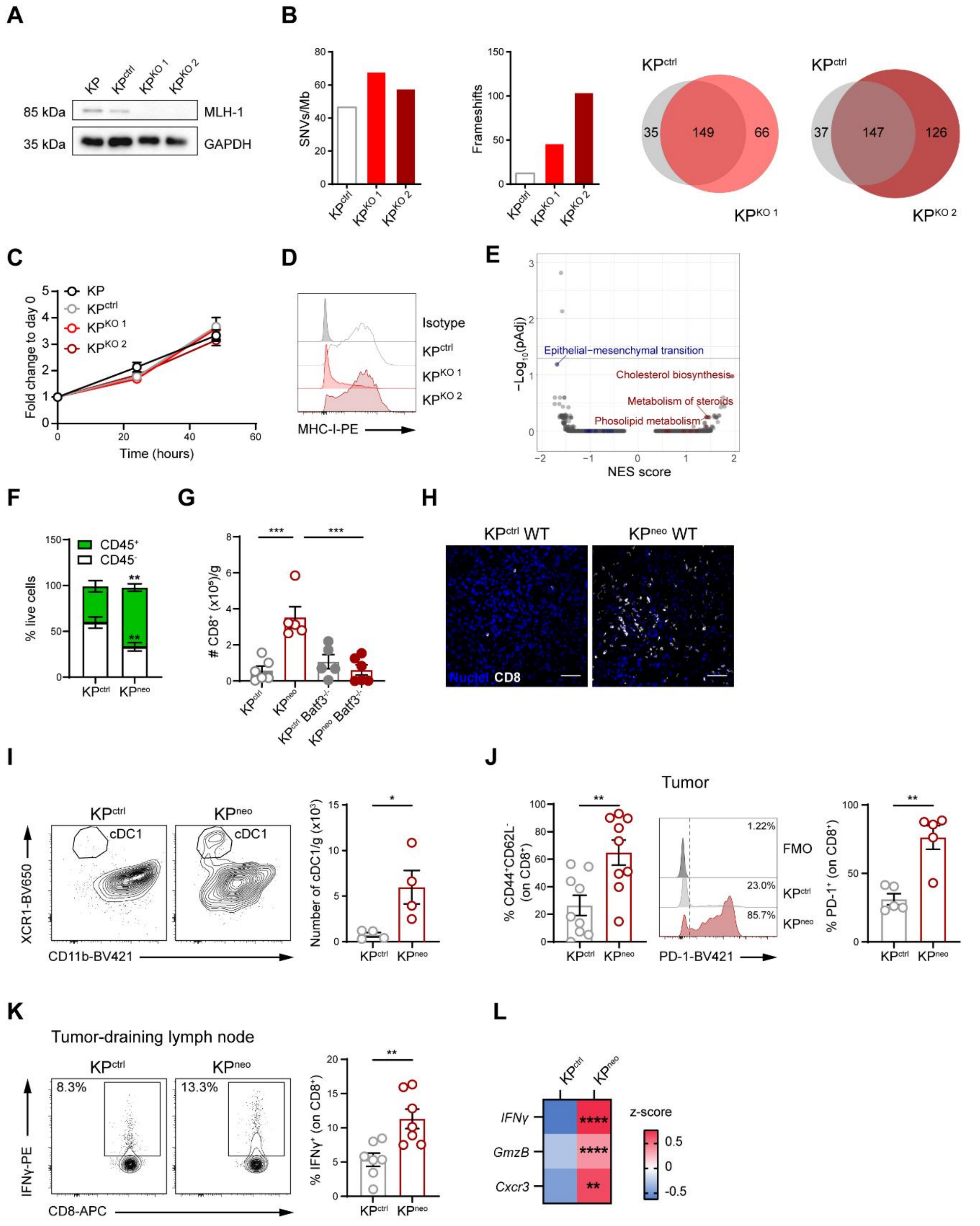


**Supplementary Figure 1. Generation and characterization of KPneo cells.**

**A)** *Mlh1* was inactivated by CRISPR-CAS9 transient transfection. KP$^{ctrl}$ cells were generated by transient CAS9 transfection without targeting vector. Cells were subcloned after transfection and screened for MLH1 expression by Western Blot. **B)** KP$^{ctrl}$ cells and two Mlh1-deficient clones (KP$^{KO1}$, KP$^{KO2}$) were sequenced (whole exome sequencing) to determine the tumor mutational load. Bars show single nucleotide variants (SNVs/Mb) (left) and frameshifts (right) in each clone. The Venn diagram depicts shared and unique predicted neoAgs calculated using NetMHCI 4.0 package and the mouse reference C57 genome as a baseline. **C)** Growth of the parental KP line and $KP^{ctrl}$, $KP^{KO1}$ and $KP^{KO2}$ was evaluated *in vitro* at 3 time points. Data refers to numbers of cells and are plotted as fold change over day 0. **D)** Levels of MHC class-I were assessed by flow cytometry (FC) after IFNγ stimulation. **E)** Gene set enrichment on RNAseq data from $KP^{neo}$ and $KP^{ctrl}$ cells showed no changes in pathways related to cell proliferation, metabolism, inflammatory responses and antigen processing (using Reactome and Hallmarks databases). NES (Normalized Enrichment Score), $KP^{neo}$ vs $KP^{ctrl}$; (adjusted p values (adj. p-value). **F-K)** $KP^{ctrl}$ and $KP^{neo}$ cells were implanted s.c. in wild type and Batf3 deficient animals. Tissues were harvested at day 21 to analyze the immune infiltrate by FC and visualize by confocal microscopy. **F)** Bars show cell fractions of $CD45^{+}$ and $CD45^{-}$ cells among live cells in tumor masses (n=9, two polled experiments). **G)** Absolute numbers of CD8 T cells infiltrating tumors in wild-type and $Batf3^{-/-}$ hosts (n=5-6, one out of two independent experiments). **H)** Representative tissue cryosections showing localization of CD8 T cells within tumor nodules. Scale bars represent 50 μm. **I)** of cDC1 infiltrating the tumor mass at day 21 post-tumor challenge (n=4, one out of two independent experiments). **J)** Expression of effector/effector memory markers ($CD44^{+}/CD62L^{-}$) and PD-1 on tumor infiltrating CD8 T cells (left n=9, two pooled experiments; right n=5, one out of two independent experiments). **K)** Tumor- draining lymph node cells (tdLN) were stimulated *ex-vivo* with PMA/Ionomycin to determine the fraction of $IFN\gamma^{+}$ $CD8^{+}$ T cells by intracellular staining (ICS). Flow cytometry analysis and quantification (n=7, pooled data form two experiments). **L)** Relative expression of the indicated genes (RT-qPCR) in $KP^{ctrl}$ and $KP^{neo}$ tumors. Heat map showing the z-score of the relative expression (RT-qPCR) of the indicated genes in total tumor tissues (n=7, pooled data from two experiments). $^{**}p<0.01$, $^{***}p<0.001$, $^{****}p<0.0001$; Two-way ANOVA followed by Tukey's test in **C**, **G**; Sidak's test in **F;** *t* test in **I,J**; and Multiple *t* tests in **K**. All data are plotted as mean ± SEM.

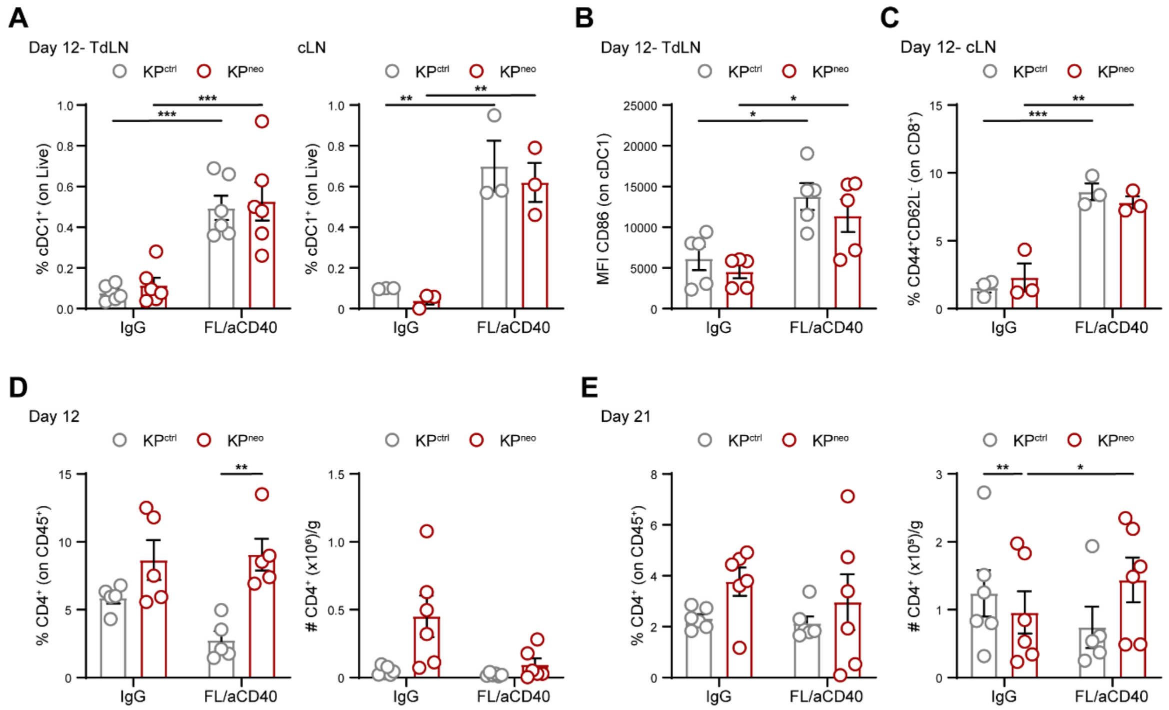

**Supplementary Figure 2. Immune profiling of DCs and CD4 T cells after DC-Therapy.**

**A)** Frequency of cDC1 in the tumor-draining lymph node (tdLN) (left) and in the contralateral non-draining lymph node (cLN) (right) at day 12 post-tumor challenge (n=3-6). **B)** CD86 expression on cDC1 in tdLN (n=5). **C)** Percentage of CD8 T cells expressing effector/effector memory markers (CD44$^{+}$/CD62L$^{-}$) in cLN at day 12 post-tumor challenge (n=3). **D,E)** Frequencies (left) and absolute numbers (right) of tumor-infiltrating CD4 T cells at day 12 (**D**) (n=5-8) and 21 (**E**) (n=6). *p<0.05, **p<0.01, ***p<0.001. Two-way ANOVA followed by Tukey's test in **A-E**. All data are plotted as mean ± SEM, and represent one out of two independent experiments.

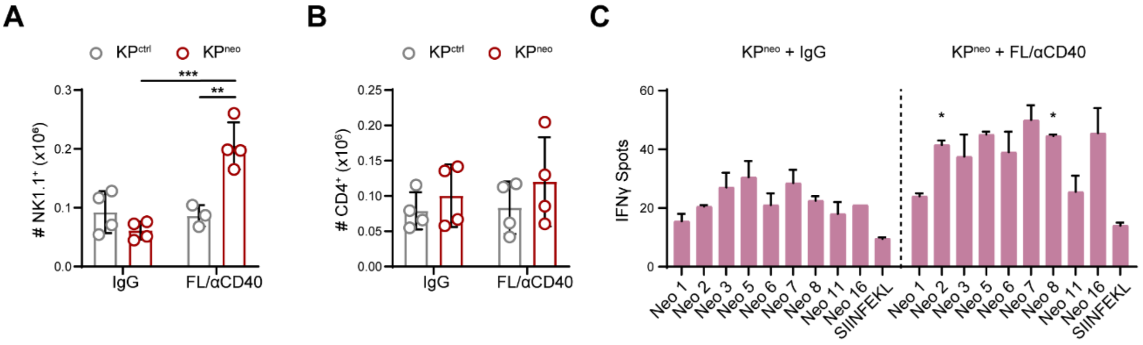


**Supplementary Figure 3. Profiling of DCs, NK cells and CD4 T cells in DC-therapy treated lungs in KPctrl and KPneo orthotopic tumor.**

Absolute numbers of NK cells (**A**) and CD4$^{+}$ T cells (**B**) in tumor-bearing lungs with KP$^{ctrl}$ or KP$^{neo}$ tumors upon FL/αCD40 or IgG, quantified by FC (n=4, one out of two independent experiments). **C)** IFN-γ ELISpot showing the specificities of CD8$^{+}$ T cells isolated from KP$^{neo}$-bearing lungs under DC-therapy or IgG to unique peptides (n=2, one out of 4 independent experiments). *p<0.05, **p<0.01, ***p<0.001. Two-way ANOVA followed by Tukey's test in **A**; Multiple *t* tests in **C**. All data are plotted as mean ± SEM.

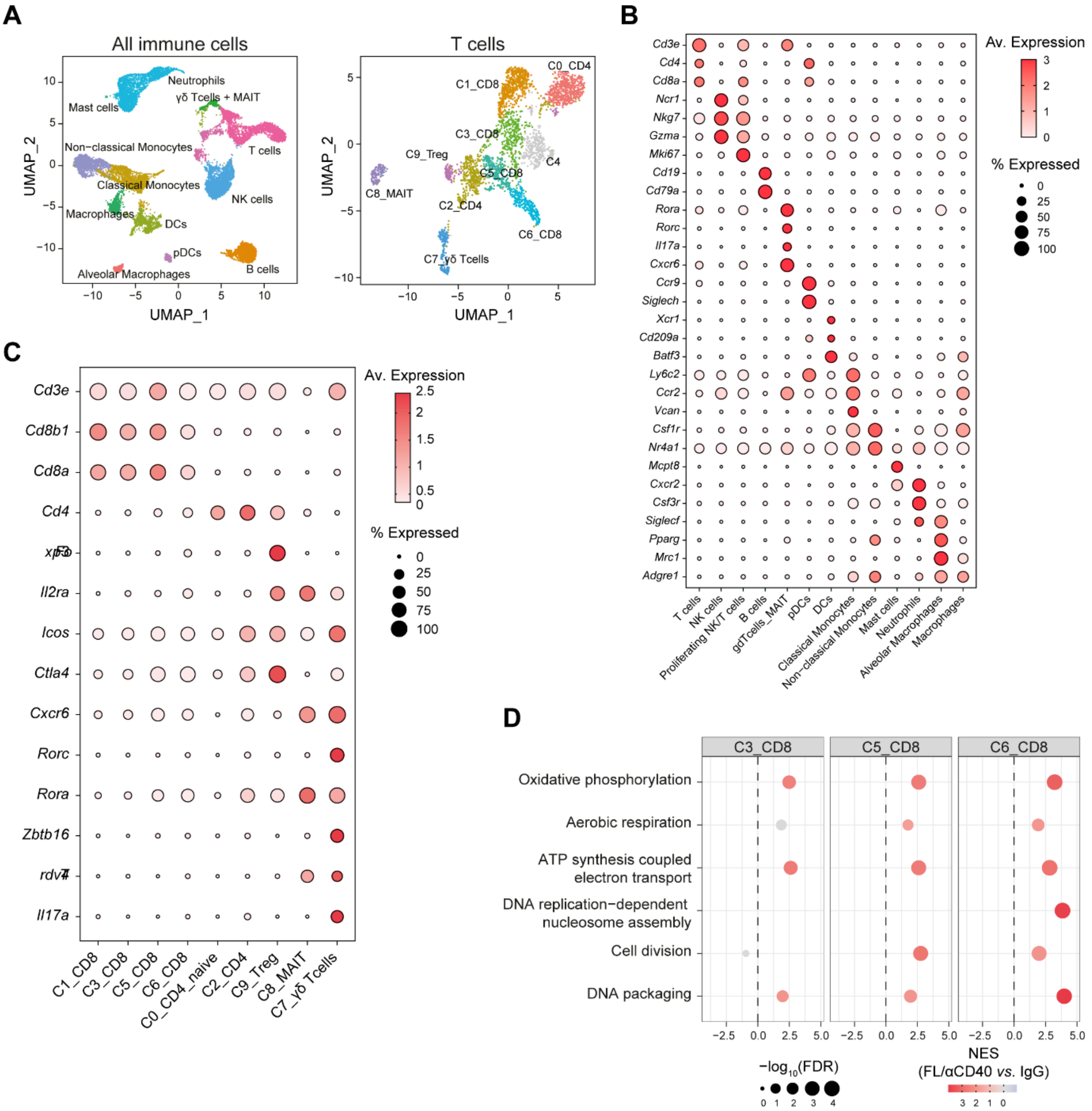


**Supplementary Figure 4. scRNA-seq of CD45+ cells from lung tissues carrying KPneo tumor upon FL/aCD40 therapy.**

**A)** CD45$^{+}$ cells were sorted from total lung tissues carrying KP$^{neo}$ tumors treated with FL/αCD40 therapy or control IgG and analyzed by scRNA-seq. UMAP visualization of CD45$^{+}$ clusters (left), and T cells clusters (right), data are merged from therapy and control samples. **B,C)** Dot plot showing expression of selected genes (negative values set to zero) defining CD45$^{+}$ clusters **(B)** and all T cells subsets **(C).** **D)** GSEA performed on expressed genes in different CD8 T cell clusters, ranked by $\log_2$FC for FL/αCD40 versus IgG comparison, using biological processes gene ontologies as gene sets. Normalized enrichment scores (NES) and significance are reported for selected significant terms.

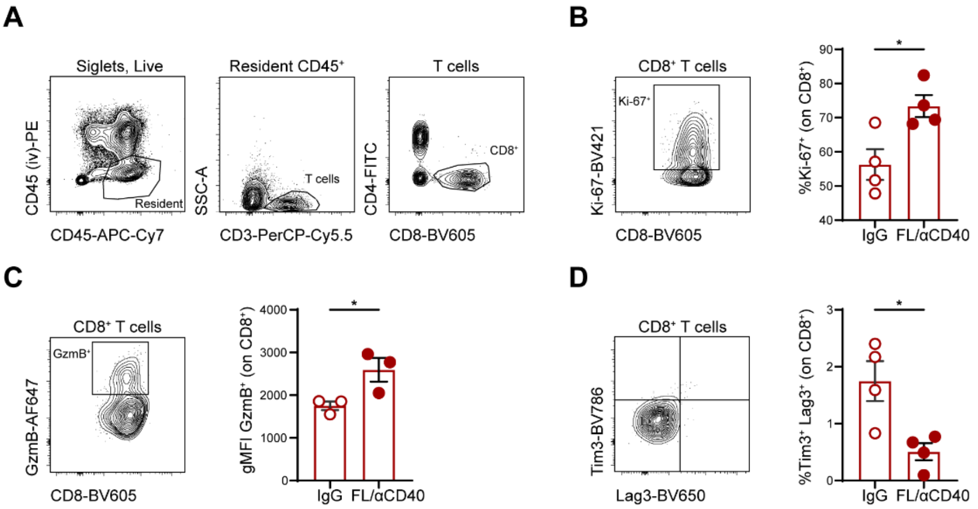


**Supplementary Figure 5. Remodeling of lung resident T cells upon FL/aCD40 therapy.**

KP$^{neo}$ tumor were implanted orthotopically in WT mice and treated with FL/αCD40 or control isotype. At the endpoint (day 9) mice mire injected intravenously with 3 μg of CD45-PE to exclude circulating immune cells. **A)** Gating strategy showing the exclusion of circulating CD45$^+$ cells and further gating to identify CD8$^+$ T cells. **B)** Dot plot and quantification of fraction of resident Ki-67$^+$ CD8$^+$ T cells (n=4). **C)** Dot plot and quantification of geometric MFI (gMFI) of GzmB in resident CD8$^+$ T cells (n=3). **D)** Dot plot and quantification of fraction of exhausted Tim3$^+$Lag3$^+$ CD8$^+$ T cells (n=4). *$p<0.05$, Student *t*-test in **B**-**D**. Data are plotted as mean ± SEM and represent one out of two independent experiments.

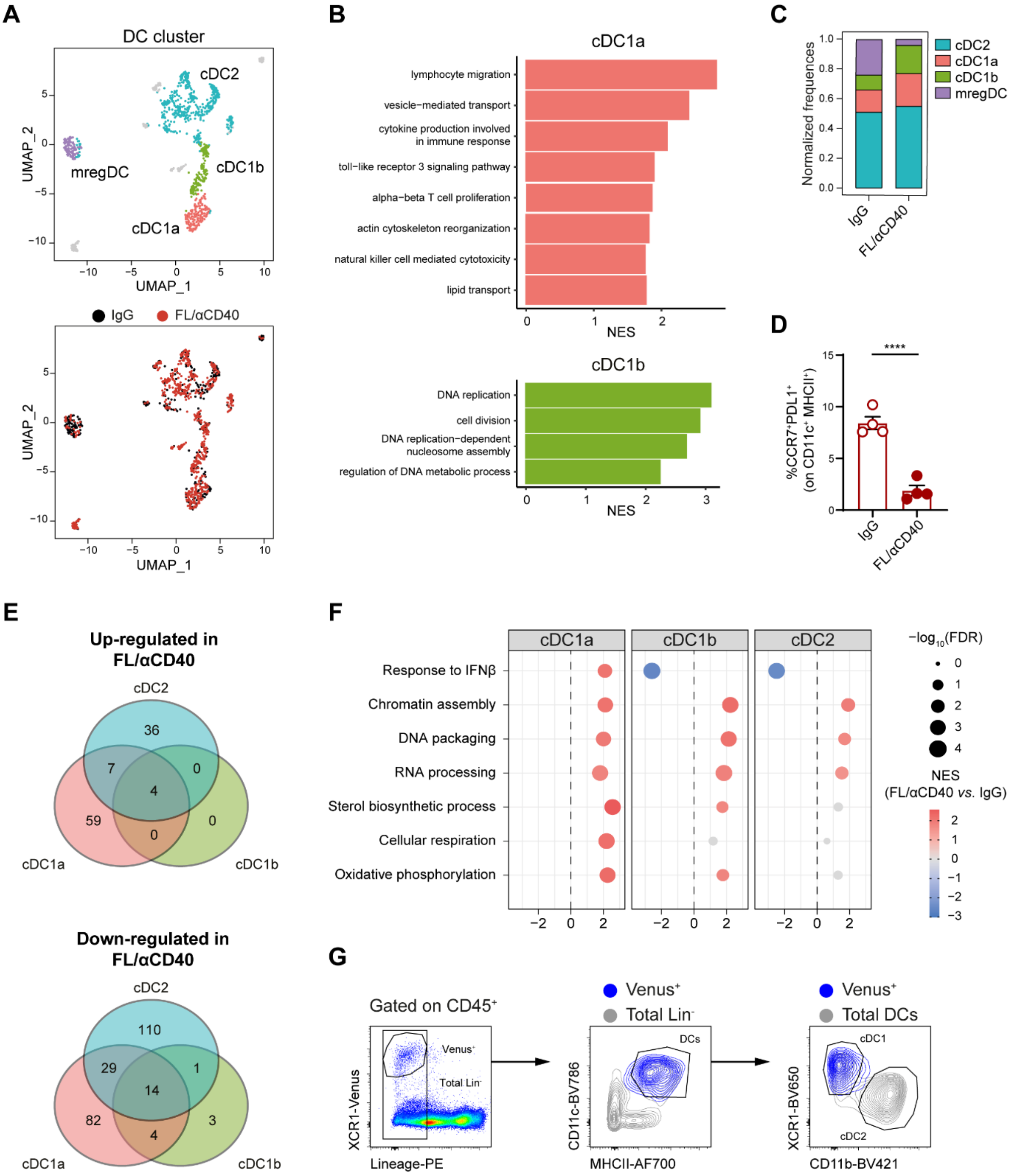


**Supplementary Figure 6. Remodelling of tissue resident cDC1 by FL/aCD40 therapy.** **A)** UMAP visualization of merged scRNA-seq data showing cells in the DCs cluster colored by sub-cluster (left), or experimental condition (right). **B)** Gene set enrichment analysis (GSEA) performed on expressed genes of cDC1s clusters ranked by $log_2$FC (cDC1a vs cDC1b comparison), using gene ontologies-biological processes as gene sets. Normalized enrichment scores (NES) are reported for selected significant terms. GSEA performed on genes ranked by $log_2$FC in cDC1a and cDC1b. **C)** Cluster composition per condition, in control (IgG) or therapy treated group (FL/αCD40). **D)** Frequencies of mregsDCs in KP$^{neo}$ tumor bearing lungs in control and therapy treated groups. mregsDCs were identified as PD-L1$^+$CCR7$^+$ on MHCII$^{high}$ CD11c$^+$ cells (n=4, one out of two independent experiments).

**E)** Venn diagrams show the number of common and unique upregulated or downregulated DEGs in therapy-treated vs control samples in the 3 main cDCs subsets. **F)** GSEA performed on all genes in each of the 3 major DCs clusters (treated vs control). NES and significance are reported. **G)** Gating strategy and representative dot plots to identify XCR1-Venus cDC1 in control and FL/αCD40 treated lung tissues. Lin (lineage) includes B220, CD3ε, CD19, F4/80, Ly6C, Ly6G and NK1.1. In gray is depicted the classical gating strategy used to identify cDC1 and in blue is overlayed the Venus$^{+}$ populations, showing the unequivocally identification of cDC1. ****p<0.0001, Student t-test in D. Data are plotted as mean ± SEM and represent one out of two independent experiments
